# Complete mitochondrial genomes of Eastern Massasauga (*Sistrurus catenatus*) and North American Racer (*Coluber constrictor*)

**DOI:** 10.1101/2024.07.24.604992

**Authors:** Ying Chen, Amy A. Chabot, Bradley J. Swanson, Stephen C. Lougheed

## Abstract

We report the complete mitochondrial genomes of two snake species of conservation concern, Eastern Massasauga (*Sistrurus catenatus*) and North American Racer (*Coluber constrictor*). The mitogenome length is 17,245bp for *S. catenatus* and 17,142bp for *C. constrictor*. Both mitogenomes include 13 protein-coding genes, 22 transfer RNA genes, two ribosomal RNA genes and two control regions. Gene arrangements are similar between two species, except that tRNA-Pro is located between tRNA-Ile and Control Region 2 in *S. catenatus*, but between tRNA-Thr and Control Region 1 in *C. constrictor*. Such gene arrangements are typical of their respective families, Viperidae and Colubridae. A phylogenetic tree with 21 additional snake species using concatenated 12 protein-coding genes (without ND6) supported current taxonomy of two reported species. We also found that North American racers in Ohio, Michigan (USA) and Pelee Island (Canada) were more closely allied to Pennsylvania individuals from the Eastern United States.

## Introduction

The Eastern Massasauga (*Sistrurus catenatus*, Rafinesque 1818) is a venomous rattlesnake in the family Viperidae (Fig. 1A). Many populations are declining and are listed as a species at risk in both the USA (U.S. Fish and Wildlife Service 2016) and Canada (COSEWIC 2012). The North American Racer (*Coluber constrictor*, Linnaeus 1758) is a non-venomous snake species in the family Colubridae (Fig. 1B). It includes 11 recognized subspecies distributed across USA, Canada, and Mexico (Integrated Taxonomic Information System, www.itis.gov). The populations in Canada are listed as endangered (COSEWIC 2002; COSEWIC 2004). Here we report the first complete mitochondrial genomes of the two species that will provide valuable resources for conservation efforts such as identifying source populations for reintroduction.

**Figure 1.**
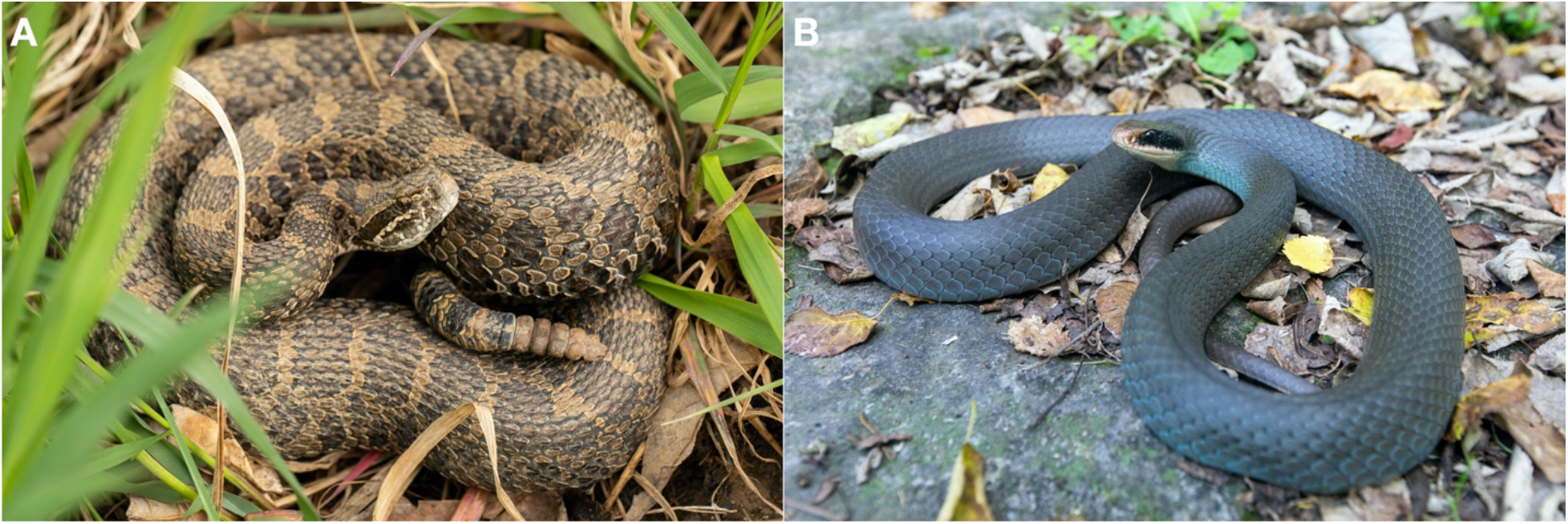
**(A)** Eastern Massasauga (*Sistrurus catenatus*) from southeast Michigan, USA. The photo was taken on May 6, 2022 by Ryan Wolfe. Eastern Massasauga is grey to brownish with saddle- or butterfly-shaped blotches on the back and alternating blotches along the sides. It has vertical pupils, triangular head with three dark stripes and a small rattle made up of interlocking segments. **(B)** North America Racer (*Coluber constrictor*) on Pelle Island, Ontario, Canada. The photo was taken on October 20, 2021 by Ryan Wolfe. North America Racer on Pelee island has pale blue or bluish green body and a whitish or bluish belly. The eyes are surrounded by a dark “mask”.

## Materials and methods

Blood was collected from a male Eastern Massasauga in Bruce Peninsula, Ontario, Canada on May 31^st^, 2009 (45.1°N, 81.4°W) and stored in 95% ethanol at - 80°C. The sample is archived in Royal Ontario Museum in Toronto, Canada (https://www.rom.on.ca/en, contact Amy Lathrop under the voucher number: ROM 50647). We extracted high molecular weight gDNA using the QIAGEN MagAttract ^®^ HMW DNA Kit (Qiagen, Germantown, MD, USA). A 10x Genomics microfluidic partitioned library (10x Genomics, Pleasanton, CA, USA) was constructed and sequenced on one Illumina HiSeqX lane at The Center for Applied Genomics (SickKids Hospital, Toronto, Canada). Each raw sequence contains a 16-base barcode that was added during the 10x Genomics library construction for the purpose of linking short reads into long contigs in nuclear genome assembly steps (Weisenfeld et al. 2017). To assemble the mitochondrial genome, we first trimmed the barcodes and primers using the python script *process_10xReads*.*py* (https://github.com/ucdavis-bioinformatics/proc10xG). We filtered out the reads marked as AMBIGOUS and UNKNOWN barcodes using *filter_10xReads*.*py* (https://github.com/ucdavis-bioinformatics/proc10xG). The filtered reads were then used to assemble the mitogenome in NOVOPlasty (version 4.3.1, Dierckxsens et al. 2016). The seed for the assembly was a GenBank sequence of *S. catenatus* spanning the NADH dehydrogenase subunit 4 (ND4), tRNA-His and tRNA-Ser region (Accession: AF156575, Parkinson et al. 2000).

For the North American Racer, blood or muscle samples were collected from 10 individuals in Michigan and Ohio in USA and Pelee Island in Canada from 2013 and 2014 (Table 1) and stored in 95% ethanol at -80°C. The samples are archived at the Royal Ontario Museum in Toronto, Canada (https://www.rom.on.ca/en, contact Amy Lathrop with voucher numbers in Table 1). The DNA was extracted using the QIAGEN MagAttract ^®^ HMW DNA Kit. Paired-end whole genome sequencing libraries were constructed and sequenced on an Illumina HiSeqX at The Center for Applied Genomics (SickKids Hospital, Toronto, Canada). The mitogenome for each sample was assembled from the raw reads in NOVOPlasty using a cytochrome *b* (cyt*b*) sequence of *C. constrictor* as the seed (GenBank accession: EU180489, Burbrink et al. 2008).

**Table 1.**
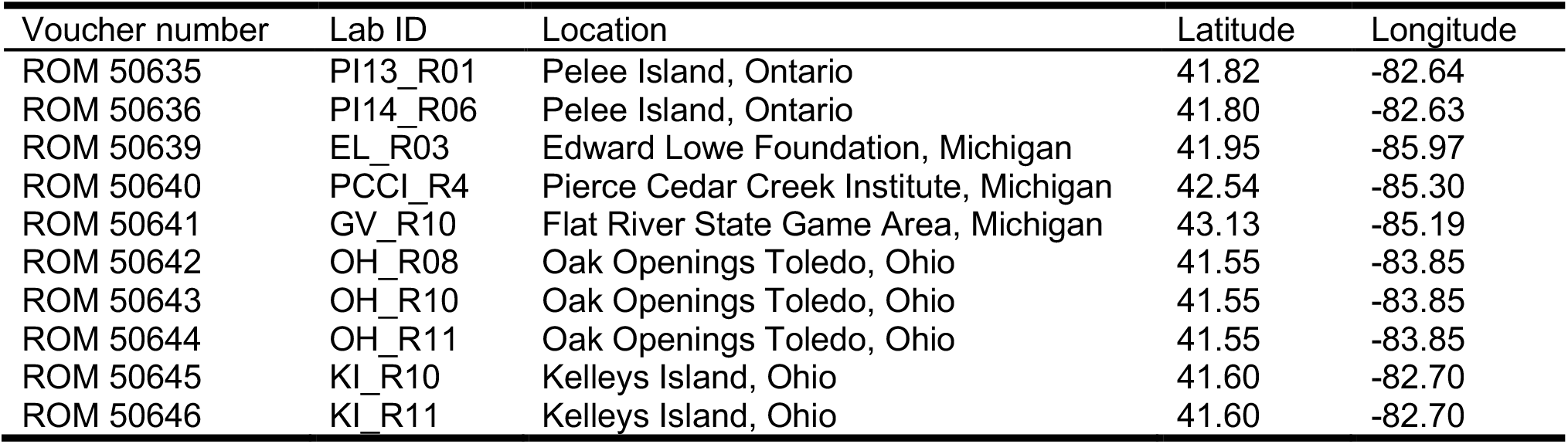
Sample locations of North America Racer (*Coluber constrictor*). The GPS coordinates are rounded to two decimal places to protect the species of conservation concern.

We annotated two mitogenomes using MITOS2 (Donath et al. 2019) and then manually checked start and stop positions of protein coding genes using blast searches (Johnson et al. 2008). The order and orientation of the genes were visualized using OGDRAW web server (https://chlorobox.mpimp-golm.mpg.de/OGDraw.html) (Greiner et al. 2019). To verify these two mitogenomes, we selected 9 members of the Viperidae, 9 of the Colubridae, 2 of the Elapidae, and 1 Xenodermidae species and downloaded their mitogenomic sequences from NCBI GenBank to construct a phylogenetic tree. We used 12 concatenated protein-coding genes, excluding ND6 as it is encoded on the light strand and thus has different base composition but also performs poorly in phylogenetic analyses (Asakawa et al. 1991; Miya and Nishida 2000). We aligned the sequences and selected the best substitution model using both the Bayesian Information Criterion and Akaike Information Criterion in MEGA X (Kumar et al. 2018). The resulting model (GTR+G+I) was used for Bayesian phylogenetic analyses in MrBayes (version 3.2.7, Ronquist et al. 2012), setting *Achalinus meiguensis* as our outgroup (GenBank accession: FJ424614, Wang et al. 2009). A total of 1,000,000 generations was run, sampling every 100, with a burnin of 25%. The final phylogenetic tree was visualized in R (Yu et al. 2017; R Core Team 2022).

We also examined the phylogenetic relationships of North America Racers from mainland Michigan and Ohio, Kelleys Island and Pelee Island (Table 1) using entire mitogenome sequences including two control regions. We aligned the sequences and determined the best substitution model in MEGA X (Kumar et al. 2018). The best model was HKY and we again ran our analyses for 1,000,000 generations in MrBayes, sampling every 100 with a burnin of 25%. We also used the cyt*b* sequences from the mitogenomes of the 10 racer samples (Table 1) and constructed a phylogenetic tree in MrBayes with an additional 22 samples from Burbrink et al. (2008) designating *Masticophis flagellum* as the outgroup (GenBank accession: MF402843, Myers et al. 2017). We aligned and trimmed the cyt*b* sequences into 1047bp in MEGA X (Kumar et al. 2018). The best substitutional mode was HKY+G. We ran the analyses in MrBayes for 1,000,000 generations and sampled every 100 with a burnin of 25%.

## Results

The complete mitogenome of the Eastern Massasauga is 17,245bp, with a base composition of 32.0% for A, 26.3% for T, 28.5% for C, 13.2% for G, and overall GC content of 41.7%. The complete mitogenome of the North America Racer is 17,142bp, with a base composition of 35.4% for A, 25.7% for T, 26.9% for C, 12.0% for G, and overall GC content of 38.9%. Both mitogenomes include 13 protein-coding genes, 22 transfer RNA genes (tRNA), two ribosomal RNA genes (12S rRNA and 16S rRNA), and two control regions (CRs) (Fig. 2). The gene orders are largely the same between the two species except for the position of tRNA-Pro. In the Eastern Massasauga the tRNA-Pro lies between tRNA-Ile and CR2, while in the North American Racer the tRNA-Pro lies between tRNA-Thr and CR1 (Fig. 2). Such gene arrangements are typical of their respective families, Viperidae and Colubridae (Yan et al. 2008; Qian et al. 2018). The Eastern Massasauga grouped with the Pygmy Rattlesnake (*S. miliarius*) in the same genus and the North American Racer clustered with all species from subfamily Colubrinae (Fig. 3), consistent with current taxonomy (Pyron et al. 2013).

**Figure 2.**
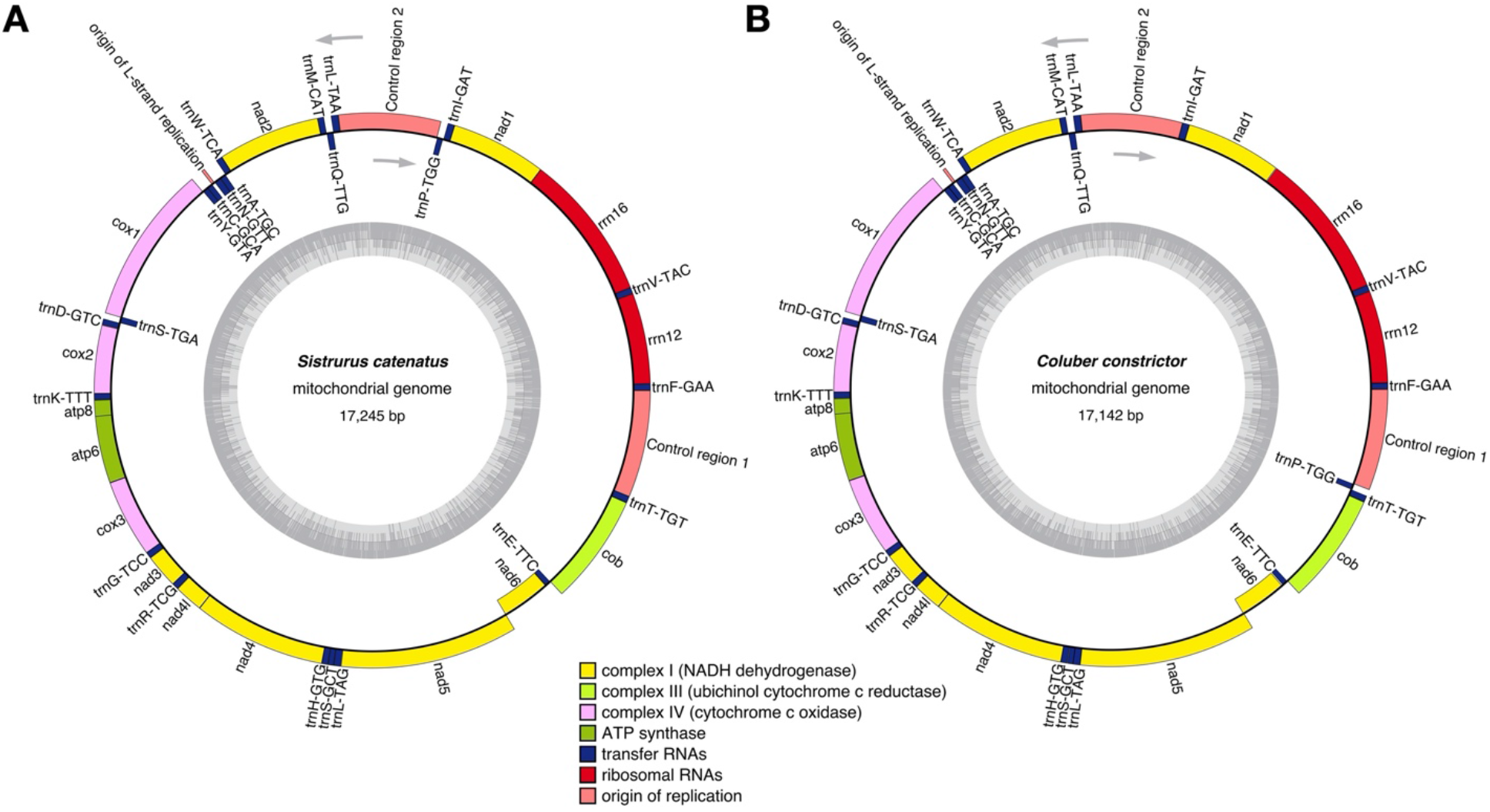
Genome map of mitochondrial genomes of **(A)** *Sistrurus catenatus* (accession OQ354766) and **(B)** *Coluber constrictor* (sample OH_R10, accession OQ354771). Both mitochondrial genomes contain 13 protein-coding genes, 22 transfer RNA genes, two ribosomal RNA genes and two control regions. The inner grey circles are the GC content.

**Figure 3.**
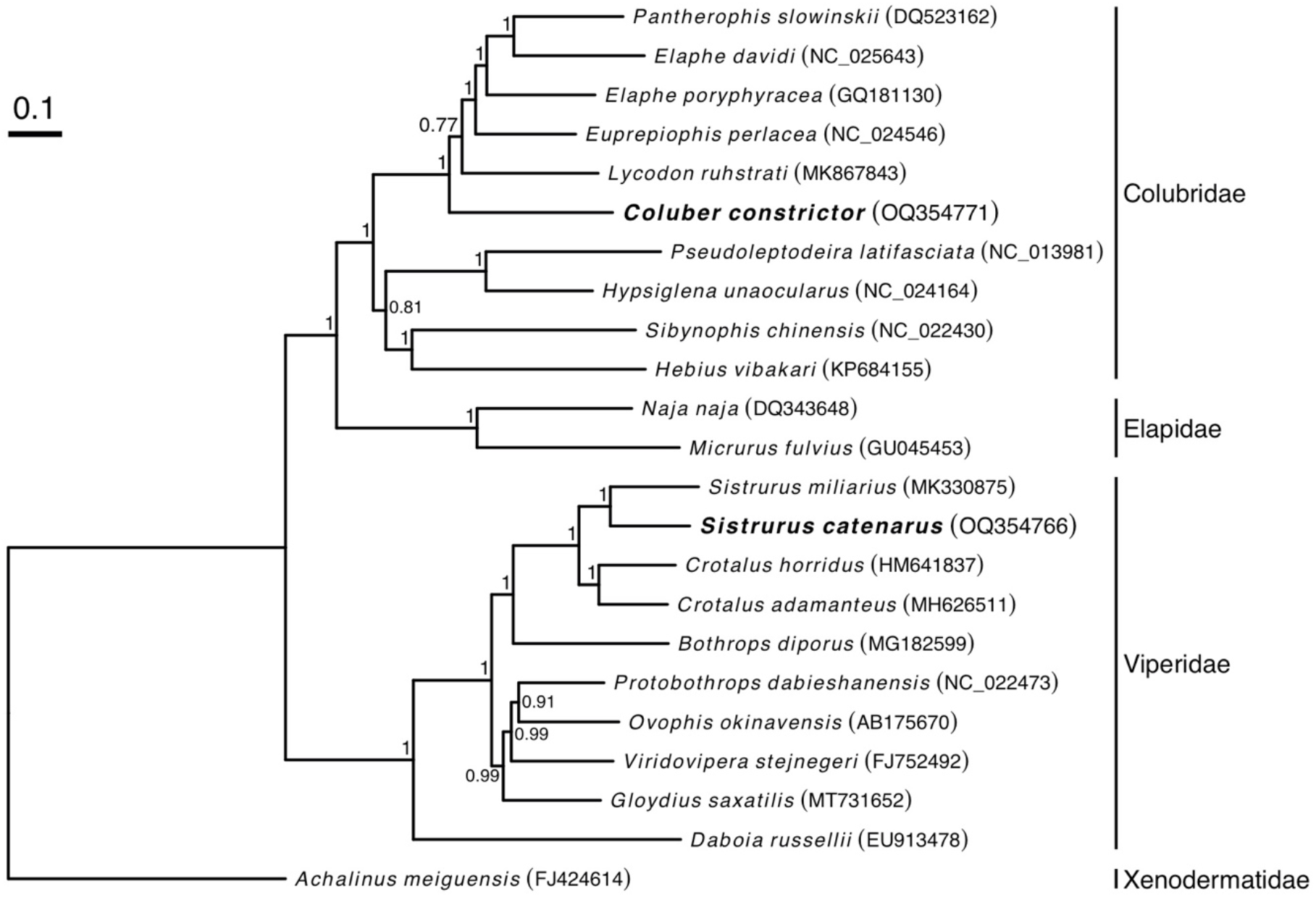
Phylogenetic relationships of 23 snake species based on 12 concatenated mitochondrial protein-coding genes (without ND6). Nodes are labelled with Bayesian posterior probabilities. The scale bar is the expected substitutions per site. The following sequences from NCBI are used in the analysis: DQ523162 (Jiang et al. 2007), NC_025643 (Xu 2014, unpublished), GQ181130 (Lin et al. 2009, unpublished), NC_024546 (Wan et al. 2013, unpublished), MK867843 (Gong et al. 2019), NC_013981 (Mulcahy and Macey 2009), NC_024164 (Mulcahy et al. 2014), NC_022430 (Oh et al. 2013), KP684155 (Xu and Zhao 2015, unpublished), DQ343648 (Yan et al. 2008), GU045453 (Castoe et al. 2009), MK330875 (Congfan 2018, unpublished), HM641837 (Hall et al. 2013), MH626511 (Wu 2019), MG182599 (Kleiz et al. 2017, unpublished), NC_022473 (Huang et al. 2014), AB175670 (Dong and Kumazawa 2005), FJ752492 (Lin et al. 2009, unpublished), MT731652 (Wu et al. 2020), EU913478 (Chen and Fu 2008, unpublished), and FJ424614 (Wang et al. 2008, unpublished).

The phylogenetic tree shows that North American Racers from Pelee Island are more related to Kelleys Island than mainland individuals (Fig. 4). North American Racers from Ohio, Michigan and Pelee Island were most closely allied with the Pennsylvania individual FTB_498 and the individuals from the Eastern United States (i.e. Eastern clade, Burbrink et al. 2008; Fig. 4), suggesting that the distribution of the Eastern clade extends farther west into Michigan (Burbrink et al. 2008).

**Figure 4.**
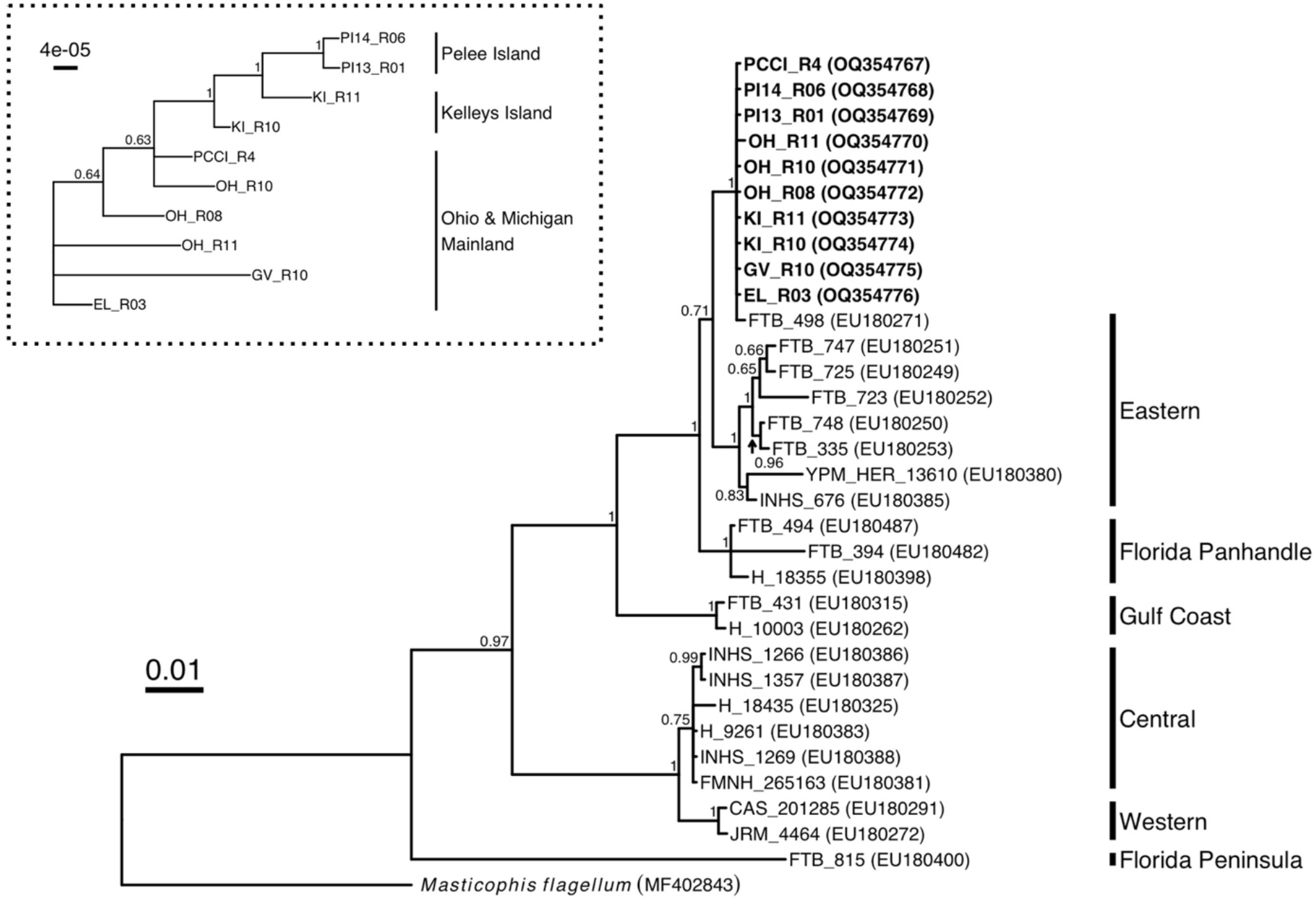
Phylogenetic tree of 10 North America Racers from this study, 22 racers from Burbrink et al. (2008) and the outgroup *Masticophis flagellum* (Myers et al. 2017) using 1,047bp of cyt*b* sequences. The phylogenetic relationships of 10 North America Racers based on the entire mitogenomes including two control regions are shown in dotted box. The labels are the mitochondrial phylogeographic lineages from Burbrink et al. (2008). Nodes are labelled with Bayesian posterior probabilities. The scale bar is the expected substitutions per site.

## Discussion and conclusion

We reported the first complete mitochondrial genomes for two snake species of conservation concern, Eastern Massasauga (*Sistrurus catenatus*) and North American Racer (*Coluber constrictor*). The phylogenetic relationships of two species are consistent with current taxonomy. We found the distribution of the Eastern mitochondrial clade of North American Racers (Burbrink et al. 2008) extends farther west into Michigan and more sampling is needed to delineate the full distribution of this Eastern mitochondrial clade.

## Ethical approval

This study is approved by the Queen’s University Animal Care Committee (reference number 2137), Canada’s Wildlife Animal Care Committee and Central Michigan University Animal Care Committee (No. 13-04A). The permits for the North America Racer are Ontario Ministry of Natural Resources #1073470, #1076138, Ontario Permit for Species Protection or Recovery AY-B-001-14, Essex Region Conservation Authority Conservation Area Use Permit No.SRACA2013-01, Michigan Department of Natural Resources WD-FRM-2014-002 316, and Ohio Division of Wildlife 14-279. The permit for the Eastern Massasauga Rattlesnake is Ontario Ministry of Natural Resources #1050742.

## Author contributions

Ying Chen extracted DNA of the Eastern Massasauga, completed all data analyses, wrote the first draft of the manuscript and revised it. Amy A. Chabot contributed to the study operations, funding acquisition, and manuscript revisions. Bradley J. Swanson contributed to the sample collection and manuscript revisions. Stephen C. Lougheed contributed to the sample collection, study operations, funding acquisition, and manuscript revisions. All authors agree to be accountable for all aspects of the work.

## Data availability statement

The sequence data produced in this study are openly available in GenBank of NCBI (https://www.ncbi.nlm.nih.gov) under the accession no. OQ354766-OQ354776. The associated BioProject, Bio-Sample numbers and SRA are PRJNA930705, SAMN33845140, SRS17116953, PRJNA946161, SAMN33816970-SAMN33816979, and SRR23922179-SRR23922188.

## Acknowledgements

We are grateful to Ryan Wolfe for providing the reference images of Eastern Massasauga and North America Racer.

